# Connectome-guided personalization of optimal tDCS intervention selection in Alzheimer’s disease

**DOI:** 10.1101/2025.08.18.670847

**Authors:** Janne J. Luppi, Annel P. Koomen, Cornelis J. Stam, Philip Scheltens, Willem de Haan

## Abstract

Transcranial direct current stimulation (tDCS) is being investigated as a clinical intervention in Alzheimer’s disease (AD) with the goal of reducing its neurophysiological effects (i.e. oscillatory slowing and loss of functional connectivity). However, progress is hampered by variable outcomes across studies, likely related to both methodological and individual differences. We recently described a virtual brain network simulation method for optimizing tDCS interventions and now propose a novel methodology for further personalizing this approach.

While the general model of the AD brain we used for our previous study was based on an average human connectome and brain anatomy, we now created new personalized models for 10 biomarker-confirmed AD patients. We used individual, amplitude envelope correlation (AEC)-based connectivity matrices extracted from magnetoencephalography (MEG) scans and implemented individual structural MRI data for current flow modeling of the tDCS effects. We then assessed a set of previously established stimulation strategies based on their ability to restore relevant neurophysiological outcome parameters in each personalized model, while undergoing AD damage.

Personalized tDCS strategies were able to delay neurophysiological deterioration, but in dissimilar ways compared to our previous results. While the general model favored posterior anodal stimulation targeting the precuneus region, the personalized models favored frontal anodal stimulation targeting the dorsolateral prefrontal cortex (dlPFC) region in 90% of the cases. This may be explained by higher connectivity levels of frontal regions in the personalized connectivity matrices, as anodal stimulation of highly connected regions produced more beneficial effects.

In this methodological study we propose several ways to improve personalized computational tDCS stimulation prediction modeling. We conclude that connectome-guided personalization of tDCS effects lead to different strategies with potentially better intervention outcomes. For external validation of this model-guided tDCS approach, personalized and general model predictions are currently being tested and compared in a clinical tDCS-MEG trial in AD patients.

## Introduction

Non-invasive brain stimulation in Alzheimer’s disease (AD) has shown promise as an alternative (or combined) treatment to pharmacological interventions [1–8]. While recent amyloid-targeting drug treatments have shown promise [9], they are expensive, intensive and not without side-effects, which is why a broad search for alternative options is justified. Transcranial direct current stimulation (tDCS) is a non-invasive brain stimulation method in which a low current is passed from a positive anode to the negative cathode in order to stimulate or inhibit the neuronal excitability of targeted regions [10–12]. The depolarizing effects of tDCS can increase neuronal activity, while hyperactivity can eventually lead to excitotoxicity (and its high metabolic demands), which is one of the established, primarily amyloid-related effects in AD. Restoration of neuronal activity within/towards a normal range can potentially lead to an improvement in neuronal network integrity, delaying the decay of cognitive processing.

The clinical appeal of this safe and patient-friendly approach is intervening directly at the level of brain function, which has a stronger relation with cognitive status than the underlying pathology [13–18]. Indirect support for effectiveness of therapy that targets neuronal dynamics comes from other neurological conditions (e.g. anti-epileptic drugs, deep brain stimulation in Parkinson’s disease), and is in line with the currently preferred AD medication restoring cholinergic deficiency, which primarily acts on neuronal signal transmission [19, 20]. Furthermore, disrupted neuronal activity can reinforce pathological protein depositions, suggesting that specific forms of restorative stimulation might be able to reduce pathological burden [21, 22]. Furthermore, non-invasive brain stimulation can promote neuroplasticity, which is highly relevant in neurodegenerative disease and not targeted with current pharmacological approaches, and can target specific regions of cortical circuits, as opposed to acting universally in the brain, enabling more diverse intervention effects, and reducing the probability of adverse events.

Despite these promising qualities, a great challenge in the field of tDCS, and non-invasive brain stimulation in general, remains the great degree of freedom in experimental design, and therefore variability in results. While several studies have found improvements in response to tDCS in both neuropsychological and clinical cognitive measures, other studies report little to no effect [23–29]. This variability is likely partly caused by differing experimental setups between studies, especially in terms of electrode placement. Often, electrode placement is motivated by directly targeting relevant cortical regions based on anatomy or pathology [8, 30, 31]. But this approach omits more distant network effects, which could, given the complex non-linear connectivity in the brain, dampen or even reverse the intended effects. Also, different stimulation regimes with regard to intensity, polarity and duration can lead to different results. This fuels the search for optimal procedures in a systematic way [29]. Secondly, there is an increasing need to personalize tDCS intervention to best suit individual anatomy, network connectivity and pathology [18, 24]. For example, certain regions may be too far atrophied in some patients to be good targets for stimulation, while other highly connected regions may predict excellent stimulation targets with network-wide intervention effects. And, with the increasing recognition of AD variants (posterior cortical atrophy, dysexecutive, frontal, corticobasal) comes the notion that these might be treated differently [32–34]. Therefore, tDCS studies could be improved by systematically justifying the electrode positions based on predicting underlying network effects in a personalized fashion [29].

How can we find a systematical approach, preferably avoiding patient burden in countless clinical trials? Computational brain network modeling is a potentially powerful option for addressing the issue of variability in tDCS and other brain stimulation research [35–38].

Neurophysiological network models can be used to run simulations in order to investigate the theoretical effectiveness of a large number of stimulation setups. This can include counterintuitive setups, which, due to the unpredictable nonlinear nature of brain networks, might still be effective [35, 37, 39–41]. Combining techniques such as current flow modeling and computational network models can be used to predict the downstream effects of the stimulation throughout networks [36, 42].

To improve the specificity of clinical model-based predictions, it is logical to consider individual variability in brain network configuration in the patient population. Differences in underlying brain anatomy or functional network connectivity may explain why a certain stimulation setup may improve cognition in certain patients with AD, but not in others [24]. For a more systematic approach to address several discussed factors of variability in tDCS literature, we have previously simulated the effects of tDCS in a neurodegenerative AD model of brain networks to theoretically optimize electrode placement in our previous work [37]. Note that while results from this previous work will inform the current study, for example in the choice of electrode montages assessed, here we will limit our analyses to the new personalized model. As before, by tuning the excitability of targeted parts of the network as a simulation of tDCS, we identified various electrode placements that resulted in improvement in neurophysiological outcome measures. Strategy success was defined by the ability to maintain or restore these measures within healthy ranges.

The aim of the current study is to build on our previous work [37], and add personalization by implementing individual anatomy and connectivity into the model to generate individualized virtual tDCS interventions. By personalizing the models used for simulating the effects of the tDCS, we determined which strategy from a previously established set of electrode montages performed best in each individual, and whether this selection differed between individuals and from the results of our past simulations in a general model without personalization. Since notable differences between individuals in structural and functional connectivity have been described, both in health and disease conditions, this might have consequences for therapy [43, 44].

We hypothesized that implementing individual anatomy and connectivity into our systematic modeling approach for optimal tDCS electrode placement in AD would result in different electrode placements performing better in certain individuals than others. As a general rule, we expected stimulation aimed at highly connected (hub) regions to perform better. External validation for the model-guided tDCS predictions will be explored in a recently completed clinical trial in which AD patients have received both general and personalized tDCS interventions with simultaneous magnetoencephalography (MEG).

## Materials and methods

For this methodological study, we selected a subgroup that was representative of our clinical trial cohort. The personalization procedure of our computational model is based on the integration of individual magnetoencephalography (MEG) and magnetic resonance imaging (MRI) data. Please refer to Figure 1 for an outline of the procedure.

**Figure 1:**
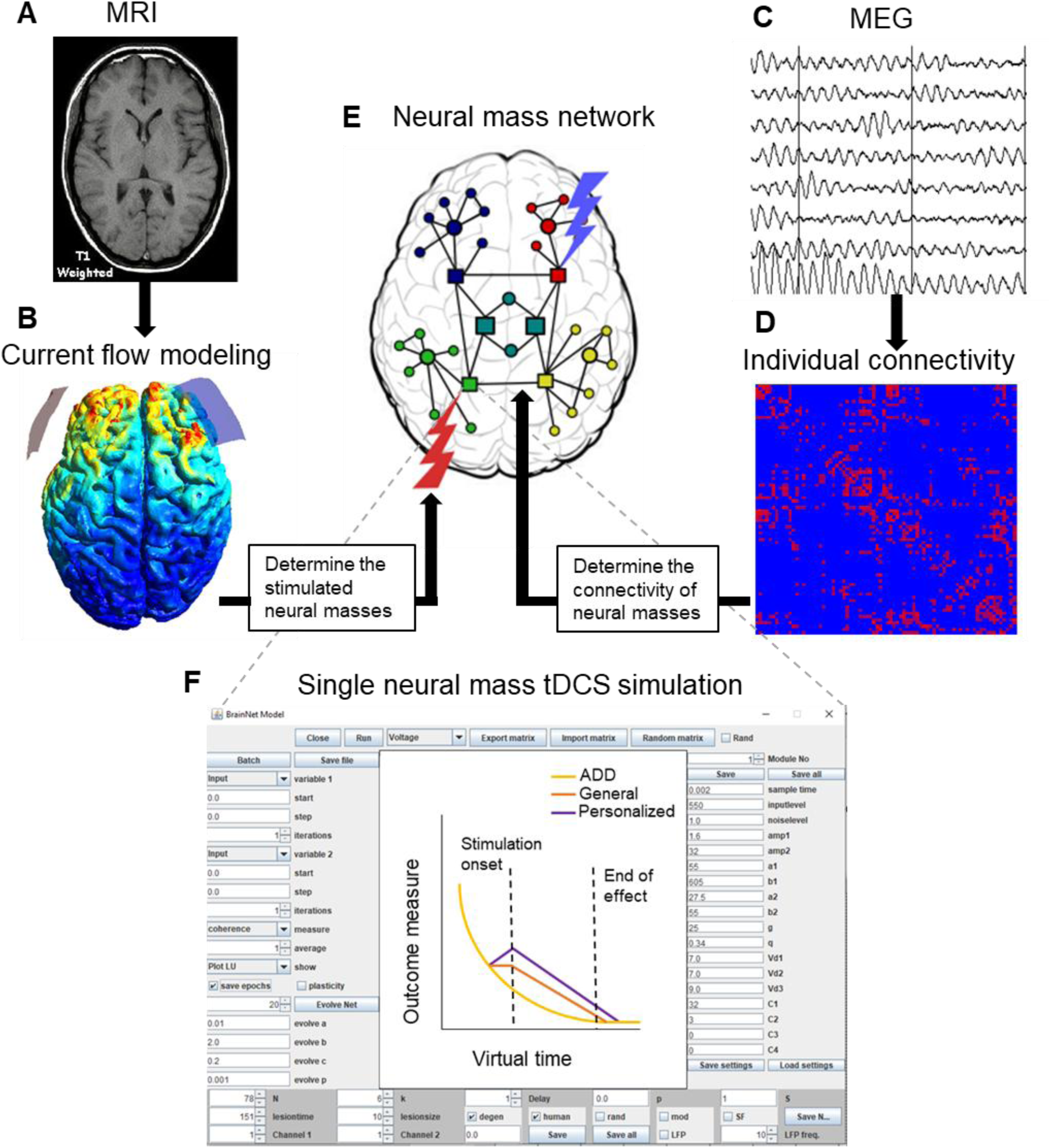
Model personalization outline. A) Individual T1-weighted images were used to generate head meshes for current flow modeling. B) Current flow modeling was used to determine which AAL atlas regions and the corresponding neural masses were stimulated in each stimulation setup. C) Individual MEG scans were used to calculate the AEC connectivity. D) Individual connectivity matrices were imported into the model. E) A schematic of the neural mass model network shows interconnected neural masses. Targeted neural masses and the connectome of the network are based on steps B and D. F) Simulations are evolved over virtual time using the activity-dependent damage algorithm (ADD) without intervention, and then again combined with all six stimulation setups. The best performing strategy in the personalized model is compared to the strategy that performed best in the general model, which is now implemented in the personalized model.

### Participants

MRI data of 10 subjects with a biomarker-confirmed early-stage AD dementia diagnosis were included for this methodological computational modeling study. Six of the participants were female, and the group had a mean age of 60.9 (± 5.9) years and an average MMSE of 23.5 (± 3.5). All participants were recruited from the Amsterdam Dementia Cohort (ADC) of the Alzheimer Center of the VU university Medical Center in the period of 2022-2024 [45]. All participants had been assessed according to a standard clinical protocol, which involved history taking, physical and neurological examination, an interview with a spouse or close family member, blood tests, 3T magnetic resonance imaging (MRI) of the brain according to a standard protocol, routine MEG, a neuropsychological assessment battery and a lumbar puncture or an amyloid-β positron emission tomography (PET). Subjects with a clinical diagnosis of dementia due to Alzheimer’s disease according to National Institute on Aging– Alzheimer’s Association (NIA-AA), and with positive amyloid-β biomarkers (cerebrospinal fluid (CSF) ptau/ amyloid-β ratio > 0.020 and/or abnormal amyloid-β PET) were recruited. The mean time between diagnosis and inclusion was 240 days (max 480 days, SD ±160 days). The local Research Ethics Committee approved the study, and all participants gave written informed consent.

### MEG data acquisition and pre-processing

Magnetoencephalography (MEG) recordings were obtained as part of the subjects’ clinical evaluation during the standard diagnostic process at the memory clinic. MEG recordings were performed in a magnetically shielded room (VacuumSchmelze GmbH, Hanua, Germany) using a 306-channel whole-head Triux Neo system (MEGIN Oy, Finland). The recording protocol consisted of a 10-minute block of eyes-closed, resting-state condition. During the scan, patients were instructed to close their eyes, stay awake, and to reduce eye movements. The recordings were sampled at 1000 Hz, with an online anti-aliasing filter (330 Hz) and high-pass filter (0.1 Hz). The temporal extension of Signal Space Separation (tSSS;[46], as implemented in MaxFilter software (Elekta Neuromag Oy, version 2.2.12), was applied with a sliding window of 10 s and correlation limit of 0.9. After visual inspection, channels containing excessive artefacts were manually removed. The head position relative to the MEG sensors was recorded continuously using the signals from five head-localization coils. The head-localization coil positions and the outline of the participant’s scalp (∼2500 points) were digitized (Fastrak, Polhemus, Colchester, VT, USA). This scalp surface was used for co-registration with individual T1-weighted MRI scans.

### MEG source reconstruction

In order to obtain source-localized activity for all regions, we applied an atlas-based beamforming approach [47]. Sensor signals were projected to an anatomical framework such that source-reconstructed neuronal activity for 78 cortical regions-of-interest (ROIs), identified by means of automated anatomical labeling (AAL), was obtained [48]. To obtain a single time series for an ROI, we used each ROI’s centroid as representative for that ROI. For the computation of the beamformer weights we used the sphere that best fitted the scalp surface as a volume conductor model and an equivalent current dipole as source model. The orientation of the dipole was chosen to maximize the beamformer output. Once the broadband (0.5–48 Hz) normalized beamformer weights for the selected voxel were computed, then the broadband (0.5–48 Hz) time-series for this voxel, i.e., a virtual electrode, was reconstructed.

### Data selection and processing

The source reconstructed time-series were converted to ASCII files for further processing. For each subject, 10 visually selected artefact-free epochs without downsampling (sample frequency 1000Hz, 4096 samples per epoch of 4.096 seconds) of the eyes-closed resting-state recording were analyzed. All quantitative spectral analyses were performed with in-house developed software (BrainWave version 0.9.164.16). Broadband power and relative power were calculated using the Fast Fourier Transform with the following bands: broadband (0.5-45 Hz), delta (0.5–4 Hz), theta (4–8 Hz), lower alpha (8–10 Hz), upper alpha (10–13 Hz), beta (13–30 Hz), and gamma (30–48 Hz). Peak frequency of the posterior dominant rhythm was calculated in the 4–13 Hz range in six posterior AAL regions in each hemisphere (superior, middle and inferior occipital gyri, calcarine region, cuneus and lingual gyrus) and then averaged to obtain a single value per epoch.

### Neural mass model

The model used is comprised of interconnected neural masses, which represent interconnected populations of excitatory and inhibitory neurons. This neural mass model is an adjusted version based on the original by Lopes da Silva et al., which has been optimized for reproducing the human alpha rhythm (8-13 Hz) [49–52]. The model has been used for several network studies on AD, due to the relevance of the alpha rhythm in characterizing the oscillatory slowing in AD [36, 37, 53]. The neural masses generate an oscillatory output comparable to MEG, relating neuronal circuit characteristics to cortical activity. For more detailed model information, please refer to previous studies [36, 37, 53].

The 78 neural masses of the model correspond to the cortical regions of the automated anatomical labeling (AAL) atlas and are coupled according to human brain topology [48]. Prior to personalization, this topology is based on diffusion tensor imaging (DTI) results by Gong et al., describing average structural connectivity of the cortex of 80 healthy adults [54]. This was the general connectivity matrix used in our previous study without personalization [37]. All connections between neural masses were excitatory and reciprocal, meaning that without external influence, the differences in activity arose from the pattern of connections.

### The activity-dependent degeneration (ADD) algorithm

To simulate damage in the network caused by AD, a previously described algorithm of activity-dependent degeneration (ADD) was used in the model [36, 53]. The ADD algorithm damages the network by lowering the coupling strength within and between neural masses as a function of recent spike density of the main excitatory population. This means that the higher the activity level in a neural mass, the more the structural connectivity of that neural mass is damaged over virtual time in the model. Since the current model has no plasticity features, i.e. no option to strengthen or regain connections, the ADD algorithm inevitably leads to network breakdown.

### Current flow modeling

Current flow modeling (CFM) was carried out in the free SimNIBS software package [42]. CFM uses MRI data to delineate tissue types within a head and assigns them corresponding conductance properties in order to predict how current spreads within the cortex for a given tDCS electrode montage. The current spread results were used at a later stage of the pipeline to determine which parts of the neural mass model would be excited or inhibited in each investigated virtual tDCS montage.

In order to create personalized head models, the *cure* pipeline provided with the software was used to transform a T1 MRI image of each subject into a head mesh used in the software. The software then used this head mesh to calculate the current spread in the cortex based on the provided electrode positions (See Supplementary Figure 1 for an example of a personal CFM result). The six electrode positions chosen for the simulations are described in Figure 2, in which three unique positions and their hemispherically mirrored versions are included [37]. These positions were chosen due to them performing the best in the previous study as well as a preliminary analysis which showed this to be the case also in the personalized model. The montage of PO8 anode and AF3 cathode performed best in the general model, and will therefore be referred to as the general strategy, although in this study it is applied in the personalized model. Simulations were carried out using 5×5 cm gel electrodes.

**Figure 2.**
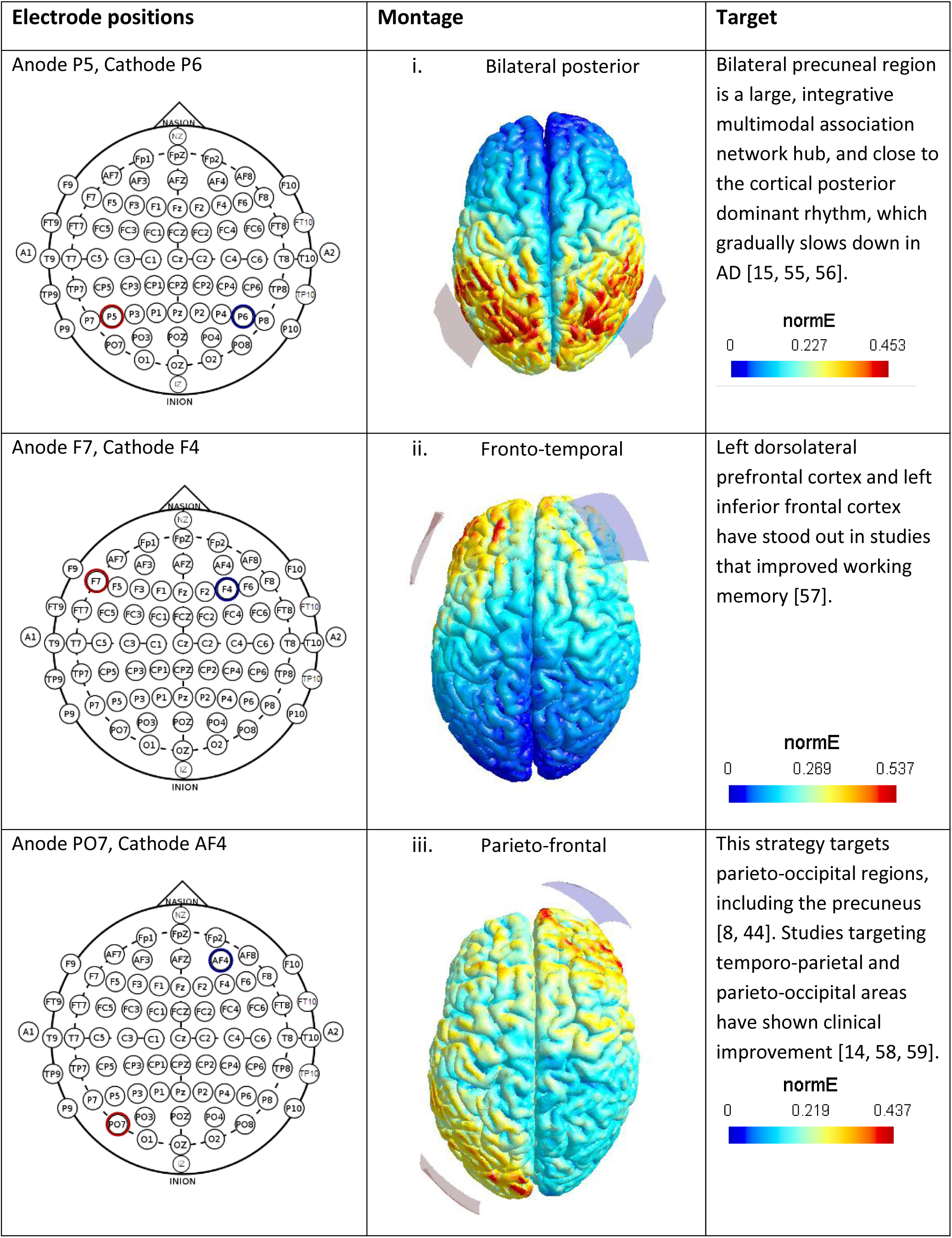
Electrode montages. The left column displays each unique electrode position used, and the middle column displays the associated current flow modeling done in SimNIBS [42]. The right column describes the intended target of interest, and the motivating literature. For each listed montage, a mirrored version was also used, e.g. in addition to an anode at F7 and a cathode at F4, a reversed position with an anode at F8 and a cathode at F3 was used.

To interpret the results, the head plots from the CFM pipeline were visually delineated into the regions of the AAL atlas, which correspond to the neural masses of the model. A region was affected by the stimulation if electric field strength exceeded 0.6 of the maximum field strength in at least 50% of region. Furthermore, each affected region was determined to be anodally stimulated if it was included in the electric field closest to the anode, or cathodally stimulated if it was included in the electric field closest to the cathode. All implementations of the CFM results into the model were carried out by the same rater.

### Individual connectivity matrices

As a measure of individual structural connectivity, the uncorrected amplitude envelope correlation (AEC) was calculated in the alpha range (8-13 Hz) based on the MEG scans of each subject [60, 61]. Because this AEC has no correction for volume conduction, it retains much of the structural connectivity information, as both are heavily influenced by distance between regions [62, 63]. It was therefore chosen as a proxy for the structural connectivity, in order for the personalized networks to be comparable to the general DTI structural network (the Gong matrix). Furthermore, since it is based on brain dynamics instead of structure, it may be more closely related to the regional activity and functional connectivity targets we are trying to restore.

The AEC was obtained for a 78×78 connectivity matrix of corresponding to the cortical regions of the Automated Anatomical Labeling (AAL) atlas. The AEC connectivity matrix was then imported into the model prior to each simulation as the personalized matrix replacing the general DTI based average human 78×78 connectivity matrix (AAL). In order to match the average connectivity of the original binarized DTI matrix with 10% sparsity while retaining the new non-binarized information, the 15% strongest connections were kept in the generated AEC connectivity matrix (see Supplementary Figure 2 for more detail). A personalized matrix averaged across the personalized matrices of all subjects was also compared to the general matrix in order to assess overlap and differences their structures.

Finally, we created an averaged AEC matrix. For this, all 10 individual AEC matrices were averaged to generate a new matrix that was then used to model strategy success in a new ‘general’ model, but now using an AEC matrix instead of a DTI matrix as was done in our previous study. This was done as a control to assess to what degree differences between the personalized models and the previous general model were caused by individual differences or by methodological differences between the AEC and DTI matrices. All simulations were performed the same on this model as for the personalized models.

### Virtual tDCS simulations

After importing the personalized connectivity matrices into the model, simulations were carried out in eight conditions for each subject. In the healthy control condition, the model was run for 30 virtual time steps with default settings, to create a baseline. In the AD damage condition, the ADD algorithm was turned on from the beginning and kept on, to simulate the AD condition without intervention. The remaining six conditions corresponded to the six virtual tDCS interventions documented in Figure 2. In each intervention condition, the virtual tDCS was turned on at virtual time point 10, after the ADD algorithm had had time to damage the network[37]. The virtual tDCS was done by changing the threshold potential of the excitatory pyramidal neurons (Vd1) of affected neural masses, as determined by the CFM results. Anodally stimulated neural masses had their Vd1 reduced from 7 to 5 (increased excitability), while cathodally stimulated neural masses had their Vd1 increased from 7 to 9. The range of these changes was based on our previous modeling work, where we found these changes to result in observable but still physiologically plausible effects [37]. The virtual tDCS was kept on for the rest on the simulation. Due to stochastic noise in the model, each stimulation step was repeated 20 times and then averaged for increased reliability and smoother visualization of the results.

### Outcome measures

As in our previous study, the same six outcome measures were used to assess spectral power (total power, relative power in the lower (8-10 Hz) and upper (10-13) alpha bands and posterior dominant peak frequency) and functional connectivity (phase lag index (PLI, [64]) and AEC in the lower alpha band, 8-10 Hz, [61, 65]). Uncorrected AEC was used, as there is no volume conduction in the model. These measures were compared between the HC and the ADD with and without intervention conditions in the personalized models.

Rather than focusing on absolute values, which can be prone to overfitting and overinterpretation, we primarily used the model to assess directions of change in response to AD-damage and virtual tDCS, which it is well suited for. An intervention was considered successful if it resulted in a shift towards the HC values in comparison to the ADD values without intervention. For example, since the ADD algorithm results in a decrease in total power in comparison to the HC condition, a successful intervention would have to cause an increase in total power compared to the ADD condition.

### Quantification of tDCS strategy success

To quantify the relationship between the degree of anodally stimulated regions and stimulation success, each personalized strategy outcome across individuals was given a measure of intervention success in the form of an area under the curve (AUC). We chose to focus on the anodally stimulated regions, because anodal and cathodal stimulation have opposing effects and in our previous study anodal stimulation was related to positive stimulation outcomes [37]. The change in all outcome measures in response to stimulation was calculated between virtual time points t=10 and t=20, as t=10 is the stimulation onset and the largest stimulation effects are seen between this and t=20. We calculated the AUC of the personalized and general strategy outcome and subtracted the AUC of the ADD outcome from this, to estimate a treatment effect. This was then repeated and averaged for all six outcome measures of each strategy, all six strategies in an individual, and all 10 individuals. This change in AUC was normalized as a Z-score to keep the influence of all outcome measures equal. Finally, the average AEC degree of each region that was anodally stimulated in at least one strategy was calculated for the virtual time point t=10 in the ADD without intervention condition. This average AEC degree was then correlated with the averaged AUC Z-score of all strategies it was anodally stimulated in. The AUC was chosen as a measure to better capture changes over time from stimulation onset, instead of focusing on a single time point.

### Hub restoration index

To assess whether virtual tDCS was able to restore connectivity to the highly connected and vulnerable hub regions, we adapted a variation of the previously described ‘hub disruption index’ (HDI) [66, 67]. This measure is calculated by correlating the baseline connectivity degree of regions in a control group with the difference in degree of the same regions in a disease group. In contrast, we reversed this measure to consider the AD damage condition without intervention as the control group and the personalized intervention condition as the comparison group. As this comparison measures the effects caused by the intervention, we labeled it the ‘hub restoration index’ (HRI), in which a higher value indicates a better treatment outcome. This index was averaged across all strategies and individuals at virtual time t=15.

### Statistical testing

Due to the model only including stochastic noise as a difference between each run of the simulations, the statistical analysis was kept simple. As mentioned, averaged measures from 20 repetitions were used as basis for all outcome measures, and these effects were further assessed in the form of the AUC changes in each condition and outcome measure.

Differences between the AUC outcomes in the ADD without intervention condition, general strategy condition and personalized strategy condition were then compared using within-subjects t-tests.

## Results

We first provide an overview of the averaged simulation outcomes in the new personalized models. We compare the best personalized strategy in each model with the previously established general strategy in the same models. Then, we further explain these results in the context of the underlying connectomes used in the new personalized and the old general models.

### Personalization leads to different tDCS predictions and performance

The personalized strategy of each model is the best performing strategy of said model, i.e. the strategy that resulted in most improvement towards healthy control values across all outcome measures. In eight cases this was F7 anode with F4 cathode, in one case F8 anode with F3 cathode and in one case it was PO8 anode with AF3 cathode, the same as the general strategy.

Figure 3 displays the results for all six outcome measures, averaged across the 10 personalized models. The personalized strategies resulted in a shift away from the AD damaged conditions and towards healthy control values in all outcome measures, therefore showing overall improvement. This change was in general present from stimulation onset at virtual time point t=10 to approximately t=20, at which point the ADD algorithm overpowered the effects of the intervention due to the lack of structural plasticity in the model. The personalized strategies outperformed the general strategies in relative lower and upper alpha power, total power, AEC and PLI, with the exception of the posterior dominant peak frequency, where the opposite was true. As such, personalizing the model resulted in a) different preferred stimulation strategies, and b) better performing strategies, although the differences were not large in the harsh ADD damage regime. The average AUC values for Figure 3 are shown in Table 1, while Figure 4 displays a representative individual case framed in the same way as the averaged results of Figure 3.

**Figure 3:**
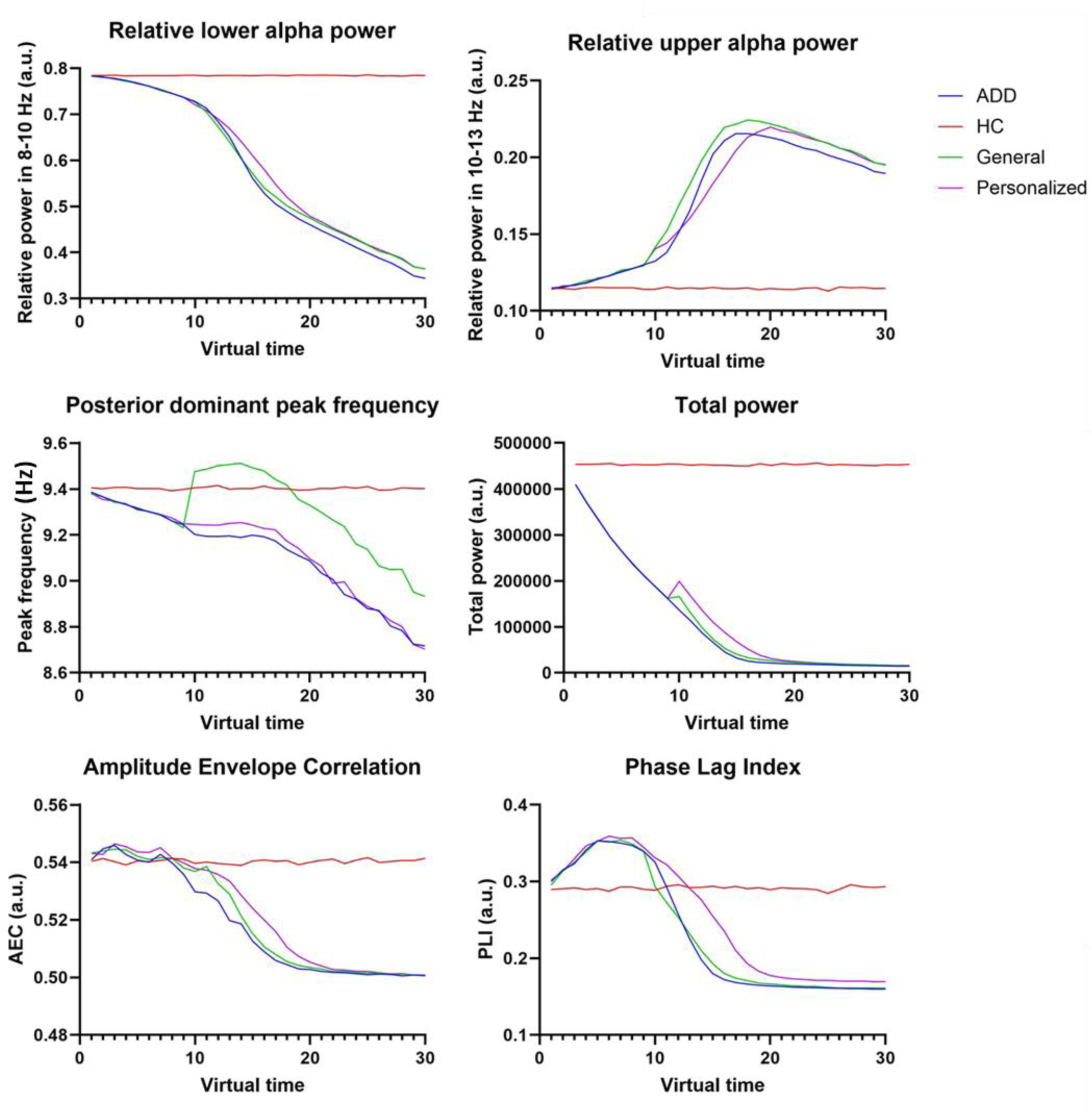
Averaged outcome measures between all individuals for personalized and general strategies. Stimulation onset was at virtual time point 10. The best performing personalized strategy and the general strategy imported into the personalized model are shown. A strategy was successful if it caused a shift from the values of the ADD condition without intervention and towards the healthy control condition. We focused on direction of change instead of absolute values, as simulations were repeated 20 times, and the variation between repetitions was small, but absolute values were also quantified in Table 1. AEC = amplitude envelope correlation. ADD = activity-dependent degeneration. HC = healthy control.

**Figure 4:**
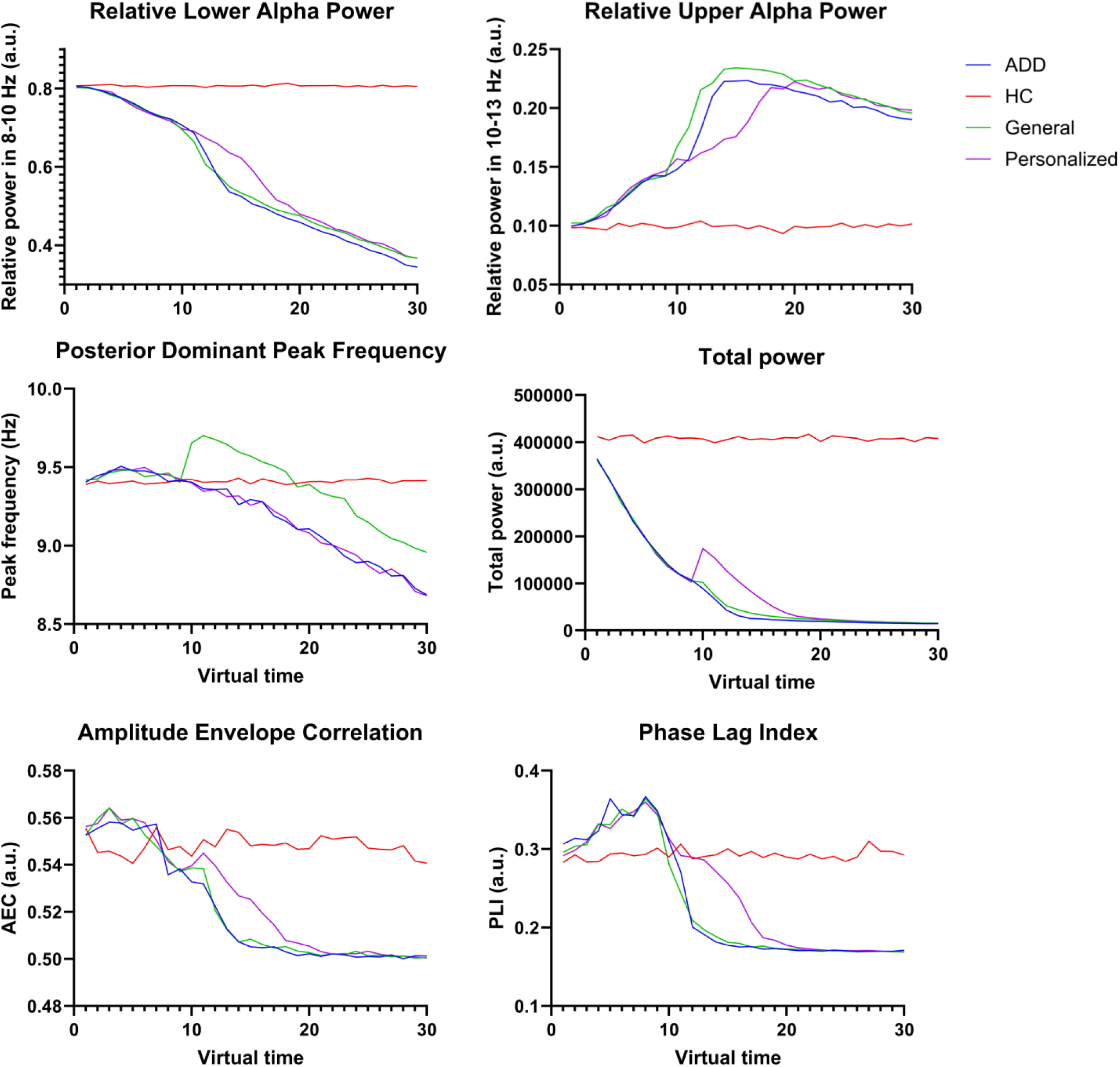
Outcome measures of a representative individual case for personalized and general strategies. Stimulation onset was at virtual time point 10. The best performing personalized strategy and the general strategy imported into the personalized model are shown. A strategy was successful if it caused a shift from the values of the ADD condition without intervention and towards the healthy control condition. We focused on direction of change instead of absolute values, as simulations were repeated 20 times, and the variation between repetitions was small, but absolute values were also quantified in Table 1. AEC = amplitude envelope correlation. ADD = activity-dependent degeneration. HC = healthy control.

**Table 1:** Effect sizes of tDCS strategies. These area-under-the-curve (AUC) values were averaged across all subjects for three conditions: the Alzheimer’s disease damage (activity-dependent degeneration, ADD) condition without intervention, the general intervention strategy and the individually best performing personalized strategy. The AUC was calculated between timepoints t=10 (stimulation onset) and t=20. Percentage change from the ADD mean is shown for in both intervention conditions, and a positive value denotes improvement (an increase, except for high alpha, for which the value was made positive to fit the others). The change in the better performing strategy is written in bold. Standard deviation is shown below each mean.

|  | Low relative<br>alpha power<br>(8-10 Hz) | High<br>relative<br>alpha power<br>(10-13 Hz) | Posterior<br>dominant peak<br>frequency | Total<br>power | AEC | PLI |
| --- | --- | --- | --- | --- | --- | --- |
| ADD without<br>intervention<br>AUC | 5,8<br>(± 0,26) | 1,9<br>(± 0,16) | 92<br>(± 0,79) | 51 x 10<sup>4</sup><br>(± 18 x 10<sup>4</sup>) | 5,14<br>(± 0,05) | 2,07<br>(± 0,35) |
| General<br>strategy<br>AUC | 5,84<br>(± 0,29)<br><u>+ 0,7 %</u> | 1,98<br>(± 0,19)<br><u>- 5,9 %</u> | 95<br>(± 0,89)<br><u>+ 3,1 %</u> | 60 x 10<sup>4</sup><br>(± 18 x 10<sup>4</sup>)<br><u>+ 17,9 %</u> | 5,19<br>(± 0,07)<br><u>+ 0,8 %</u> | 2,08<br>(± 0,32)<br><u>+ 0,6 %</u> |
| Best<br>personal<br>strategy AUC | 6,06<br>(± 0,27)<br><u>+ 4,5 %</u> | 1,82<br>(± 0,15)<br><u>+ 2,8 %</u> | 92,17<br>(± 0,64)<br><u>+ 0,5 %</u> | 82 x 10<sup>4</sup><br>(± 23 x 10<sup>4</sup>)<br><u>+ 62,4 %</u> | 5,23<br>(± 0,05)<br><u>+ 1,7 %</u> | 2,53<br>(± 0,33)<br><u>+ 22,1 %</u> |

### Connectome source (DTI vs MEG) choice influences tDCS performance

Due to the difference in best performing stimulation strategies in the personalized and the general models, the differences in their connectivity were investigated. Figure 5 displays the overlap between the connectivity matrices based on the general DTI and the average of the personalized AEC results. While the matrices have a similar overall structure, one key difference is that the AEC matrix features more strong frontal connections (please see the green frontal clusters in figure 5).

**Figure 5:**
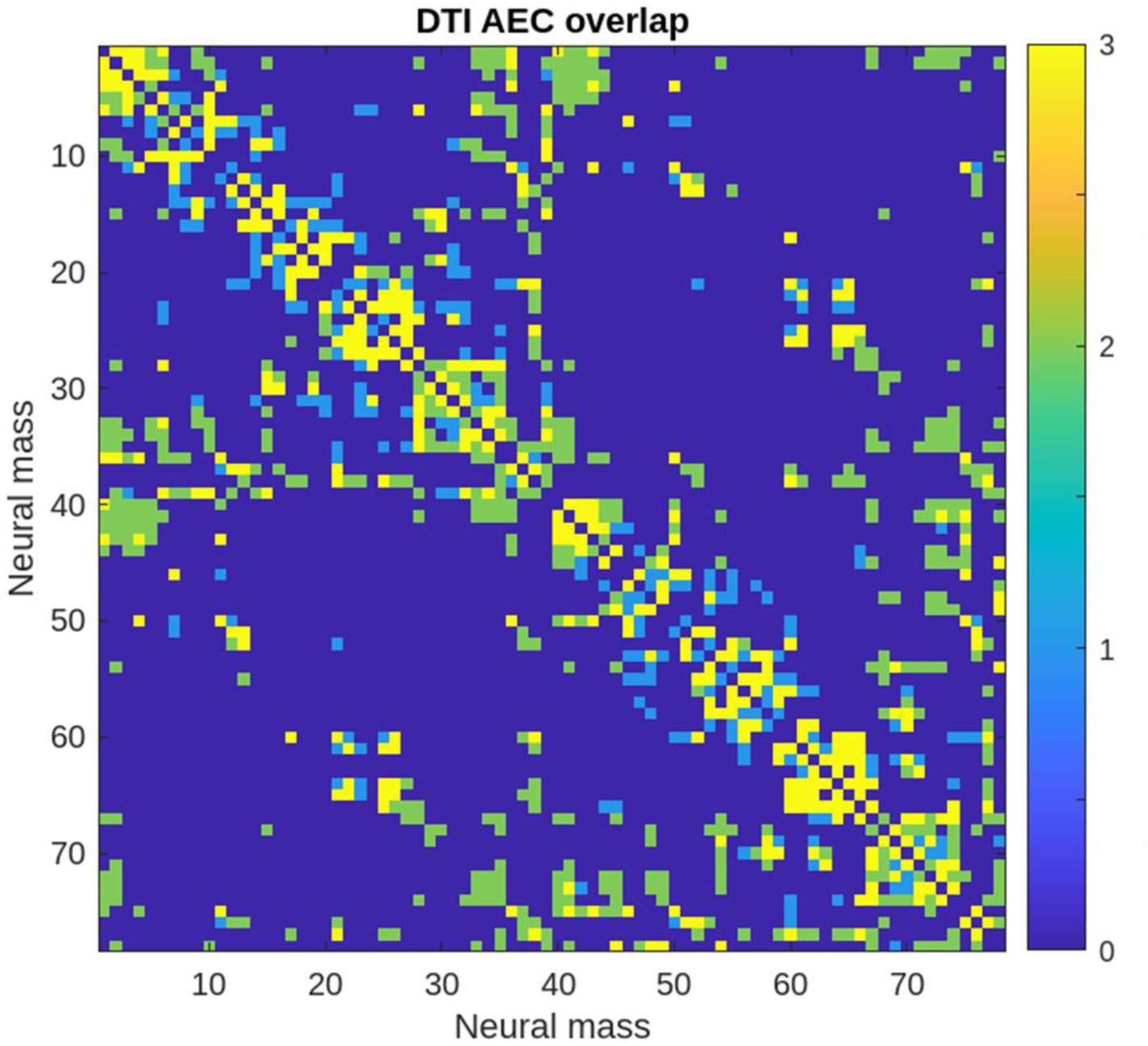
Overlap between the original DTI-based connectivity matrix and averaged personalized AEC-based connectivity matrix. A dark blue square denotes a lack of connection in both matrices, a light blue point denotes a connection only in the DTI matrix, a green square denotes a connection only in the AEC matrix, while a yellow point denotes a connection shared by both matrices.

### Comparison with a model with an averaged AEC matrix

Figure 6 displays results in all outcome measures over 10 epochs from the stimulation onset in an alternative model in which an averaged matrix calculated from the AEC of all 10 subjects was used. These figures were generated for 10 virtual time points from the stimulation onset (time point 10 in Figure 3). While there are many similarities to Figure 3, for example in that the F7 anode with F4 cathode montage performed best and the relationships between conditions being identical for total power, differences are also present. Unlike in Figure 3, posterior dominant peak frequency does not favor the personalized strategy as clearly, and AEC favors the general strategy.

**Figure 6:**
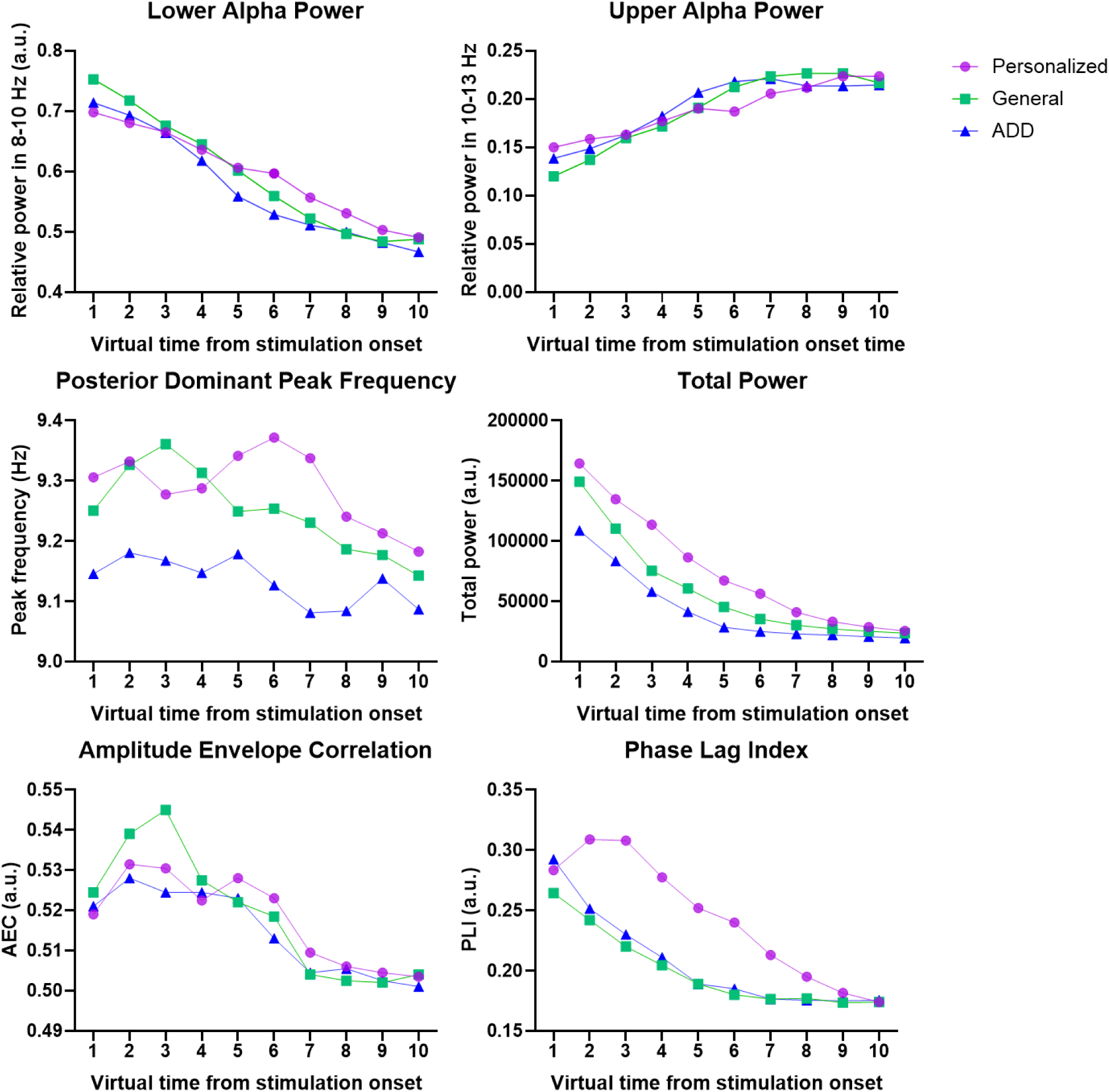
Outcome measures using an averaged AEC matrix from 10 subjects. The best performing personalized strategy (F7 anode with F4 cathode, which also performed best in this model) and the general strategy imported into the personalized model (PO8 anode with AF3c cathode) are shown over the first 10 virtual time epochs after stimulation onset. AEC = amplitude envelope correlation. ADD = activity-dependent degeneration.

*Network topography influences tDCS performance*Next, we quantified the relationship between stimulation success and the averaged degree centrality (AEC, 8-10 Hz) of anodally stimulated regions, in the form of an averaged AUC change. As seen in Figure 7, significant, moderately strong positive Pearson’s correlation was observed (r=0.434, p<0.001), indicating that anodally stimulating higher degree regions resulted in a greater intervention success.

**Figure 7:**
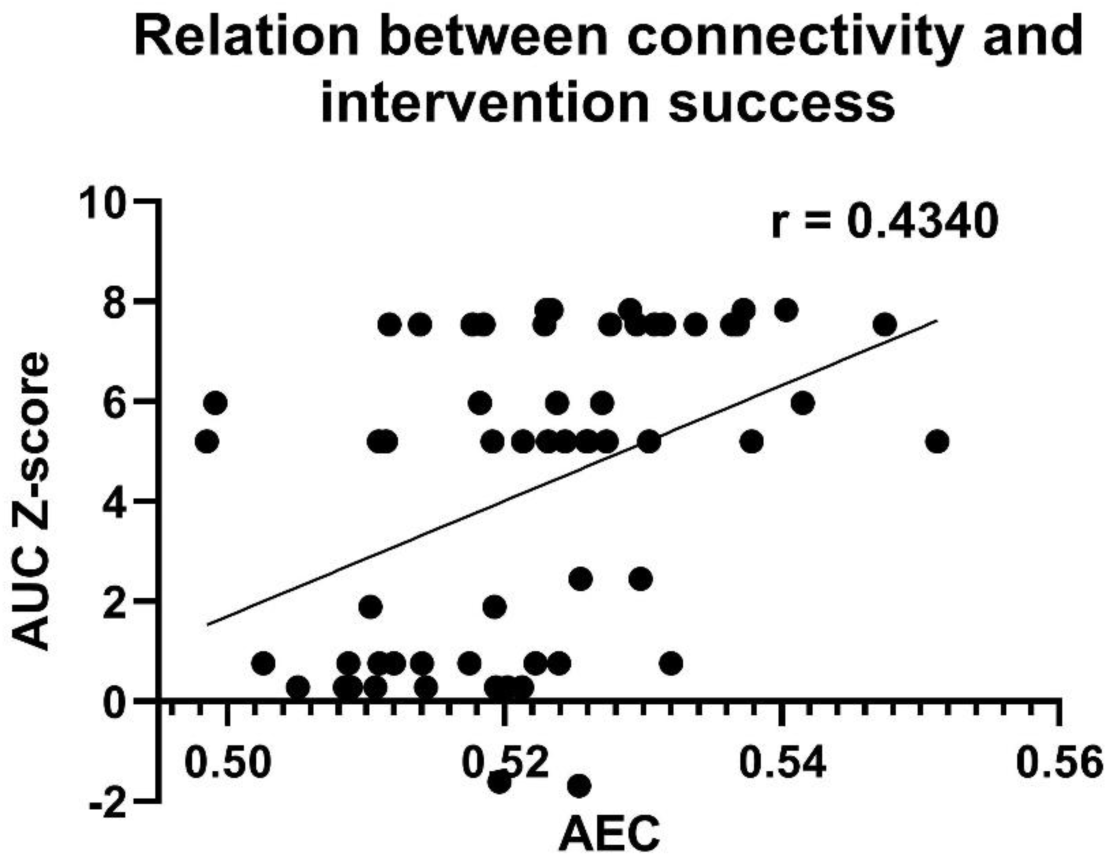
Correlation between the AEC degree of anodally stimulated regions and the AUC Z-score as a measure of personalized intervention success. Each personalized strategy of was given a measure of intervention success in the form of an averaged area under the curve (AUC) change across all outcome measures in those strategies between virtual time points t=10 and t=20, which was normalized as a Z-score. The average AEC degree of each region that was anodally stimulated in one or more of the personalized strategies was correlated with the average AUC Z-score in those strategies. A significant, moderately strong positive Pearson’s correlation was observed (r=0.434, p<0.001).

Table 2 displays the ranking of anodally stimulated regions by degree for both the personalized strategy (F7 anode, F4 cathode) and the general strategy (PO8 anode, AF3 cathode), in a representative individual case. The average AEC degree prior to stimulation was significantly higher in the areas that were anodally stimulated in the personalized strategy (p < 0.0001).

**Table 2:** Ranking of the anodally stimulated regions by degree. For a representative individual case, the regions that were anodally stimulated in the personalized strategy (F7 anode, F4 cathode) and the general strategy (PO8 anode, AF3 cathode) were ranked by degree as measure by AEC (amplitude envelope correlation). Degree was measured at virtual time t=9, prior to stimulation onset. L = left hemisphere, R = right hemisphere. The mean degree was significantly higher for the regions anodally stimulated in the personalized strategy.

| **Anodally stimulated neural masses in personalized strategy** | **Associated AEC degree** | **Anodally stimulated neural masses in general strategy** | **Associated AEC degree** | **P-value Mann-Whitney** |
| --- | --- | --- | --- | --- |
| Middle temporal pole gyrus L | 0.581 | Rolandic Operculum R | 0.585 |  |
| Superior temporal pole gyrus L | 0.577 | Heschl gyrus R | 0.568 |  |
| Rolandic operculum L | 0.574 | Superior temporal gyrus R | 0.539 |  |
| Heschl gyrus L | 0.569 | Middle temporal gyrus R | 0.536 |  |
| Rectus L | 0.569 | Postcentral gyrus R | 0.530 |  |
| Temporal superior gyrus L | 0.562 | Superior occipital gyrus R | 0.527 |  |
| Frontal medial orbital gyrus L | 0.561 | Supramarginal gyrus R | 0.524 |  |
| Frontal inferior orbital gyrus L | 0.559 | Lingual gyrus R | 0.524 |  |
| Frontal inferior triangulate gyrus L | 0.559 | Cuneus R | 0.521 |  |
| Frontal superior orbital gyrus L | 0.556 | Precuneus R | 0.519 |  |
| Frontal middle orbital gyrus L | 0.545 | Inferior parietal gyrus R | 0.518 |  |
| Frontal superior medial gyrus L | 0.541 | Parietal superior gyrus R | 0.514 |  |
| Postcentral gyrus L | 0.540 | Angular gyrus R | 0.511 |  |
| Inferior temporal gyrus L | 0.539 | Calcarine fissure R | 0.511 |  |
| Insula L | 0.539 | Middle occipital gyrus R | 0.508 |  |
| Frontal superior gyrus L | 0.529 | Inferior occipital gyrus R | 0.504 |  |
| Mean | 0.556 | Mean | 0.527 | <0,0001 |

Figure 8 displays the hub restoration index, which had a strong, significant slope of 0.7836 (p<0.0001), indicating that the hub regions with a higher AEC degree without intervention responded to anodal stimulation with the highest increase in AEC degree.

**Figure 8:**
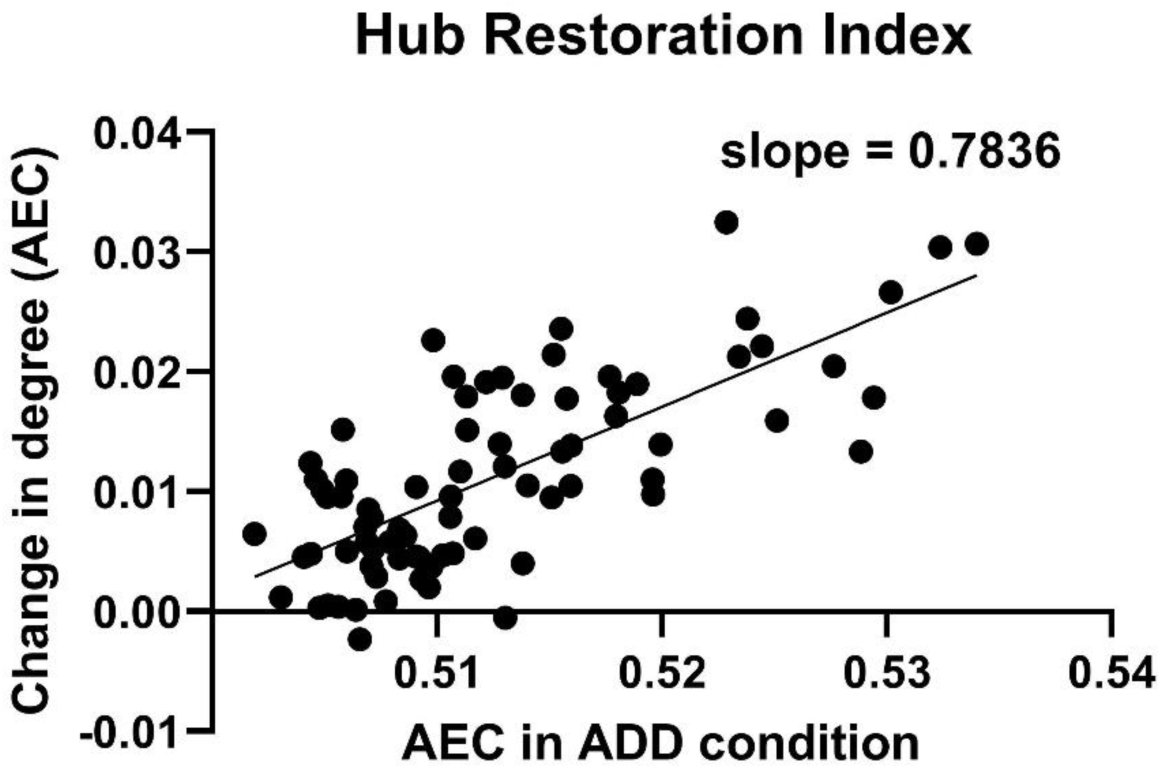
Hub restoration index for the anodally stimulated regions. To assess the topological consequences of stimulation, a hub restoration index was calculated. For each region, its AEC degree in the control condition was correlated with the comparative average change in AEC degree in all personalized interventions it was anodally stimulated (both taken at virtual time point t=15). A significant, strong slope of 0.7836 (p<0.0001) was found, indicating that anodally stimulating high degree regions resulted in a stronger restoration of AEC connectivity.

## Discussion

In this study we aimed to explore the influence of implementing individual brain structure and connectivity in a previously published computational model simulating tDCS treatment in AD, to improve the prediction of tDCS treatment response. The stimulation effects of the personalized strategies resulted in improvement in all outcome measures compared to the ‘no intervention’ condition, and in all outcome measures except the posterior dominant peak frequency compared to the general strategy condition. Personalizing the model resulted in a dlPFC-targeting electrode setup performing best in all but one individual (for whom the general strategy performed best), contrasting the results of the general model, which favored a precuneus-targeting montage. Favorable treatment response was correlated with anodal stimulation of highly connected hub regions and appeared to be especially restorative for the functional connectivity of these same regions. Here, we discuss the implications of our findings, their relation to existing literature, limitations and potential future directions.

### Model personalization leads to different tDCS strategies

We found that personalizing the neural mass model based on individual AEC-based network connectivity resulted in a different strategy performing best in nine out of ten individuals, compared to our previous results in the non-personalized model. Instead of the previously most successful general strategy (with a posterior anode near the precuneus (PO8) and a contralateral frontal cathode (AF3)), a strategy with a temporofrontal anode (F7 or F8) and a frontal cathode (F4 or F3) performed best. These frontal strategies involve the dorsolateral prefrontal cortex (dlPFC), which is one of the popular targets for tDCS interventions due to its involvement in the default mode network [13, 68–70]. These strategies were also successful in the general model, but were outperformed by the precuneus targeting montages in effect size [37]. Furthermore, targeting the dlPFC has been effective in tDCS treatments of depression [71]. Overall, these results indicate that introducing personal network connectivity notably affects the predictions, but the consistency within the personalized group might also imply that the primary cause of the observed difference is the choice for functional data instead of structural, and not the individual connectivity details [23, 24].

When comparing the connectivity matrices of the personalized and the general models, certain features stood out as differences. The overall structure of the matrices was quite similar, as can be appreciated in Figure 5. However, the sparsity of the matrices was 10% for the general and 15% for the personalized matrices. This was done to match the average connectivity degree between the two, since measures can be dependent on network size and density. The general matrix was binarized between values of 0 and 1, while the personalized matrix values were kept continuous in order to retain information, with average values for active connections being closer to 0.5 than 1. Therefore, more connections needed to be included in the personalized matrix to balance network density by adjusting the threshold to include the strongest 15% of the connections, but this may have had other unanticipated effects on the results. Moreover, the personalized matrices included topological differences, such as more frontal connectivity clusters than the general matrix. By investigating the connections with the highest degree, it was also confirmed that the connections present in the personalized but not the general matrix were not simply weaker connections that were introduced as the matrix was made less sparse. This presence of higher frontal connection density is relevant, as it could explain why frontal stimulation strategies performed better in the personalized model. While the connectivity matrices might also be different because they are derived from different modalities, we decided to accept this difference, because ultimately, we are interested in finding out whether adding different types of empirical data (i.e. MRI, MEG) helps restoring functional network integrity, regardless of dissimilarities. Of course, whether the observed difference between these strategies will be similar and equally effective in the clinical experiments is yet unknown, but from the future discrepancies between the virtual and clinical results we hope to understand which methodological choices are most valid.

### Connectivity matrix differences: driven by population or methodology?

There are three key differences in how the connectivity matrices of the personalized and general models were made: the personalization itself, as well as the population and the connectivity measure and modality the matrices are based on. Firstly, the personalized model aims to be more suited to the individual, and therefore could lead to different and potentially more accurate predictions of intervention outcome. This perspective is difficult to validate before the upcoming clinical trial comparing the efficacy of the personalized and general models in predicting optimal stimulation strategies. However, the personalization could already have an effect on the results through differences in the population used for the matrices. The general human DTI matrix was based on data form healthy adults, whereas the personalized AEC matrices are based on 55-80 old patients with a biomarker-confirmed AD diagnosis. Therefore, the personalization process could be integrating features of AD pathology and aging that were not present in the general matrix. Interestingly, some studies report increases in frontal connectivity that are theorized to be a form of compensation and predict less severe AD progression [72], while others claim that such increases predict pathology [73]. Therefore, incorporating such features into the model via personalization could benefit both prediction accuracy and our understanding of regional connectivity changes in AD.

Another possible explanation for the observed differences could be caused by the different connectivity measures used. In the general model, a purely structural connectivity measure was used, while the personalized model uses uncorrected AEC as a proxy for structural connectivity. This is because while a corrected AEC is a functional connectivity measure, leaving out the volume conduction correction results in a matrix more closely related to structural connectivity. Naturally, this can result in differences between the MRI-based DTI and the MEG-based AEC, as well as the remaining influence of functional connectivity on AEC. Moreover, uncorrected AEC is predisposed to finding spurious, false-positive connections in the midline of the brain. The reliability for the MEG beamformer reconstruction of deep sources is smaller, especially near the midline, and nearby virtual channels look more similar than they should. This can lead to spurious connections when using uncorrected FC measures, and could be behind at least some of the frontal connections that were observed in the AEC matrices but not in the DTI matrix.

To address this question we additionally ran simulations in a model based on an averaged AEC matrix across our study population, as shown in Figure 6. This served as a comparison to see whether personalized models would still have different results when compared to an averaged connectivity model based on the same AEC methodology. As could be expected, results from this model were similar to the results of the personalized models (Figure 3), but differences were also present. The best performing strategy, as in 90% of the personalized models, was the F7 anode F4 cathode strategy, and many of the relationships between the conditions were the same. For example, total power was highest in the personalized strategy, second-highest in the general strategy, and lowest in the condition without intervention. However, while posterior dominant peak frequency was highest in Figure 3 in the general strategy, in the new averaged AEC model it was highest in the personalized strategy. Additionally, AEC results in Figure 3 were highest in the personalized strategy, while in the average AEC model they were highest in the general strategy. As such, while these results are not conclusive and would benefit from a larger sample size, the differences observed indicate that differences between the personalized and general model outcomes are not based solely on the different methodologies used.

### Hub status of stimulated regions correlates with intervention success

A moderately strong significant correlation was found between the degree of anodally stimulated regions and their involvement in successful personalized strategies. In essence, the more highly connected regions were more likely to be anodally stimulated in the best performing strategies. This was also observed in the finding that the average degree of anodally stimulated regions was higher for the personalized strategy than the general strategy in a representative individual case. Again, this could support the explanation that the higher frontal connectivity in the personalized model compared to the general model resulted in more frontal strategies performing better. It could be that stimulating the less damaged frontal regions is more effective in spreading the beneficial effects of the tDCS intervention throughout the network. This may be even more important in individuals with more advanced AD pathology, as the connectivity of the personalized models was based on AD patients instead of healthy adults. Interestingly, we performed the same comparison in our previous publication on the general model, and found no such correlation [37]. This remains the case even when repeating our updated methodology of using AUC scores on the previous dataset. It is not clear what is behind this difference, but it could be related to the aforementioned differences in population and connectivity measure. Intuitively, targeting highly connected hubs appears like a sound strategy to improve controllability, and recent studies using tDCS in chronic pain have shown support for this approach [74, 75]. However, whether this is true in the case of the current model and Alzheimer’s disease requires upcoming clinical validation, especially given the possibility of them being influenced by the aforementioned midline artefacts.

### Restoration of hub connectivity

Selective hub vulnerability is a well-established phenomenon in Alzheimer’s disease, making hub regions a potential therapeutic target. Recent studies exploring the role of disrupted hub connectivity have used the ‘hub disruption index’ (HDI) as a tool for investigating the selective global and regional vulnerability of highly connected brain network regions in AD [66, 67]. This concept is closely tied to the damage caused by the ADD algorithm in our model, as damage based on activity level affects hubs more severely than less connected regions [44, 53]. To counter this, network control theory supports targeting hubs for maximal network influence [76, 77]. We therefore introduced a reversed version, the ‘hub restoration index’, to assess whether our interventions could help restore hub connectivity. We found a strong positive slope of high significance, with anodal stimulation of higher degree regions resulting not only in connectivity improvement, but also restoring neuronal dynamics relatively effectively. This lends support to the personalized interventions being able to intervene at the level of a key neurophysiological feature of AD. Naturally, this is only one potential way in which network health can be improved. For example, interest in intervening at the level of excitation-inhibition (E/I) balance, to reduce metabolic demand and optimize neuronal firing response repertoires, has been increasing in previous years, due in part to its potential in attenuating the metabolically high demands of hub hyperexcitability [67, 78, 79].

### Limitations and future directions

Naturally, the methodologies used in this experimental new approach to personalizing tDCS in AD are not without limitations. Firstly, although the focus is on individual results, the study could benefit from a higher number of participants included and strategies tested. For this personalization study, we opted for a modest number of strategies and individuals, as increasing their number causes a multiplication of the already considerable number of separate simulations and analyses needed. However, the strategies tested were selected from the 6 out of 20 strategies that performed best in the general model, and during preliminary analyses these 6 strategies also performed the best in the personalized models. Secondly, the translation of the current flow modeling results to the neural mass model changes involves laborious fitting of the CFM results to the AAL atlas used in the model. Both of the issues could possibly be alleviated if some of the processes could be further automated.

For example, instead of working with fixed electrode montages, some CFM software solutions allow for optimizing montages to target specific regions. This approach could help limit the number of montages that need to be simulated and analyzed per participant, while simultaneously improving the accuracy of the targeting. This could be further improved by choosing another atlas as a basis for the model, preferably with more uniform regions shape and size than the AAL atlas, leading to an easier and more accurate conversion of the CFM results into model settings.

Further, the difference in study population and connectivity measure of the personalized and general matrices make it challenging to compare and explain the results between the two. This was a necessity caused by a lack of access to DTI data and the need to include AD patients and not healthy adults in the study. This comparison could be made more straightforward by generating a new connectivity matrix based on AEC or another connectivity measure in a larger sample of the AD population, and then using this as a basis for the general model. While we addressed this issue with and average AEC matrix based on our study population, these differences must be held in mind when interpreting our results.

Finally, while the assumption that restoring network function will lead to clinical improvement in cognition is logical, it has not yet been proven. Ultimately, our predictions will naturally need to be validated in the upcoming clinical trial comparing MEG measurements during personalized and general tDCS interventions, which can also help inform us about optimal methodology for further personalization of the interventions.

While the BrainWave software used in this study is made available for free use, we acknowledge that going towards a more systematic and unified use of toolboxes for analysis across research groups would be beneficial. For example, non-invasive brain stimulation toolboxes such as BEST [80] and MEG analysis toolboxes such as FLUX [81] could help in bringing results such as these to other groups in a more standardized form. BrainWave was used due to our groups familiarity with it, but more widely established options hold merit for future works.

Finally, it is good to keep in mind that the present study focuses on short term neuromodulation effects, while we know that most clinical gain is to be expected with repeated, longer stimulation regimes. Neuroplasticity, which might be one of the main mechanisms to counter neurodegeneration, is not taken into consideration in this work. This is because adding plasticity greatly complicates a model and can easily make the network unstable, although in an ideal case its inclusion would make the model more realistic. As such, instantaneous, direct neuromodulatory effects of tDCS were the focus of this study, as they will be in the clinical validation phase in which we will record the effects of ongoing tDCS on AD patients’ brain activity as measured via MEG. We believe that, given the neuronal hyperexcitability in AD, beneficial changes can arise from the neuromodulatory effect of tDCS, which in turn can lead to longer-term changes via plasticity.

## Conclusion

We have described a novel methodological approach for integrating individual brain anatomy and connectivity into a neural mass model-based approach of optimizing tDCS interventions in AD for further personalization. We observed that the personalized approach resulted in a different, more frontal electrode montage being favored over a more posterior montage previously indicated as the best in a general, non-personalized model. These differences appear to be related to the connectivity of the anodally stimulated regions and the outcomes demonstrate potential in restoring connectivity of damaged hubs. General and personalized virtual tDCS model predictions will be validated in an ongoing sham-controlled clinical tDCS-MEG trial, in which MEG recordings during and immediately following general and personalized strategies will be compared.

## Acknowledgements

Research of Alzheimer center Amsterdam is part of the neurodegeneration research program of Amsterdam Neuroscience. Alzheimer Center Amsterdam is supported by Stichting Alzheimer Nederland and Stichting Steun Alzheimercentrum Amsterdam. The clinical database structure was developed with funding from Stichting Dioraphte. WH is a ZonMw Memorabel (733050518), Alzheimer Nederland (WE.03-2022-08), Hersenstichting grant recipient and ZonMw TOP (40-00812-9817043) and Open Competition (09120012110032) grant co-recipient. This studty was funded by the ZonMw Memorabel grant.

## Supplementary material

### Incorporating individual brain anatomy into the model

The free SimNIBS current flow modeling software was used to create individual head meshes based on T1 MRI images prior to predicting current spread [42]. Supplementary Figure 1 shows an example case. In most cases, personalization did not result in great differences in areas considered to be stimulated, unless considerable atrophy was present.

**Supplementary Figure 1:**
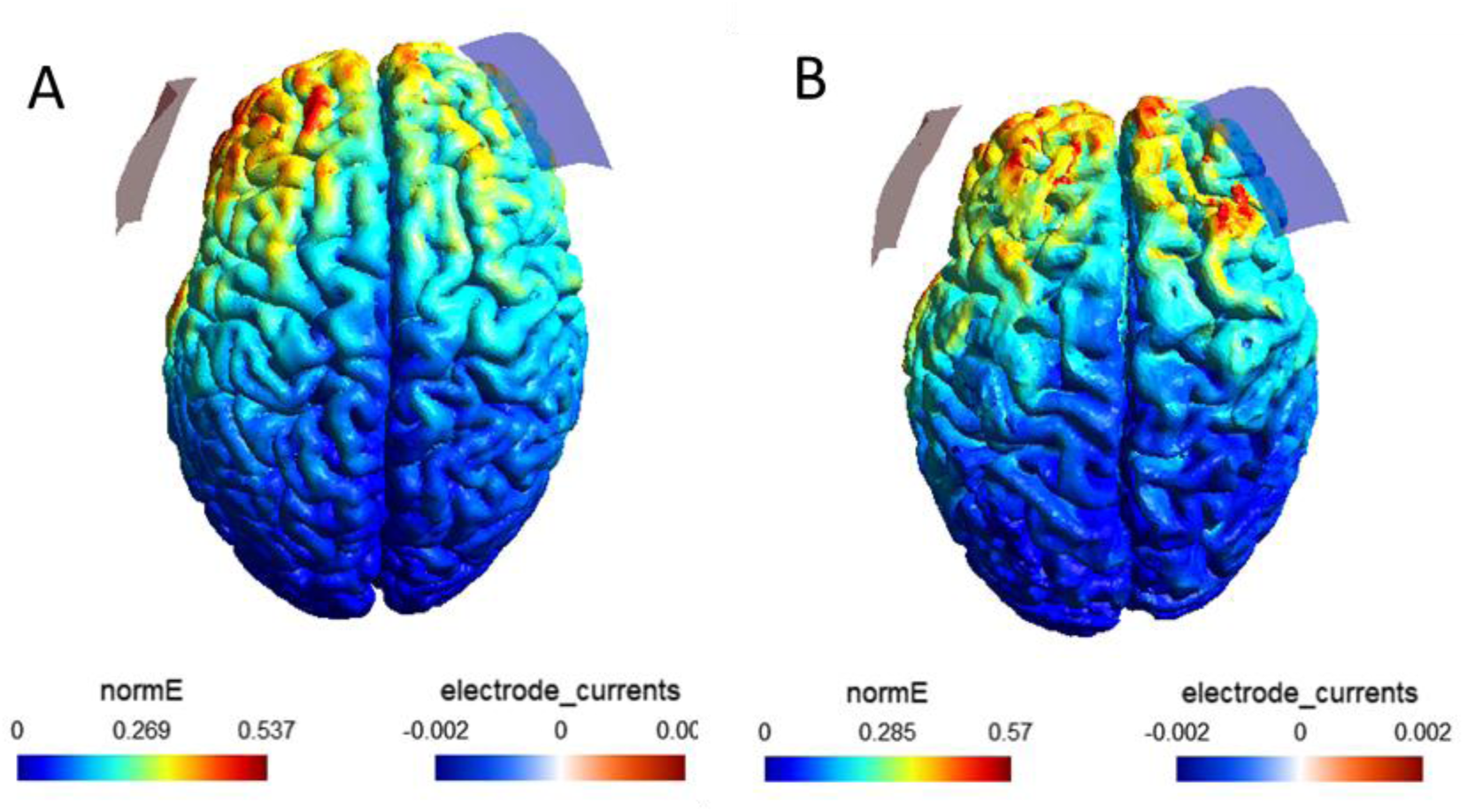
Current flow modeling on a personalized head mesh. Individual T1-weighted MRI images were used to create personalized head meshes for current flow modeling (CFM) in SimNIBS software [42]. A) A template head mesh of a healthy adult brain (Ernie) provided with the software. B) A personalized head mesh of an individual Alzheimer’s disease patient. The head mesh was visually divided into the 78 cortical regions of the AAL atlas, and regions in which at least half of the region was receiving an current field of 0.6 of the maximum strength (normE) were considered to be stimulated.

### Determining the connectivity sparsity of the personalized matrices

To balance out the fact that the personalized matrices were not binarized like the general diffusion tensor imaging matrix from Gong et al. [54], they were made less sparse in order to reach a more comparable baseline level of connectivity between the matrices. To compare model behavior at different sparsities, it was run without the damage algorithm for 20 steps and three outcome measures were analyzed, as seen in Supplementary Figure 2. The 15% sparsity threshold was chosen due to its similarity to the general model.

**Supplementary Figure 2:**
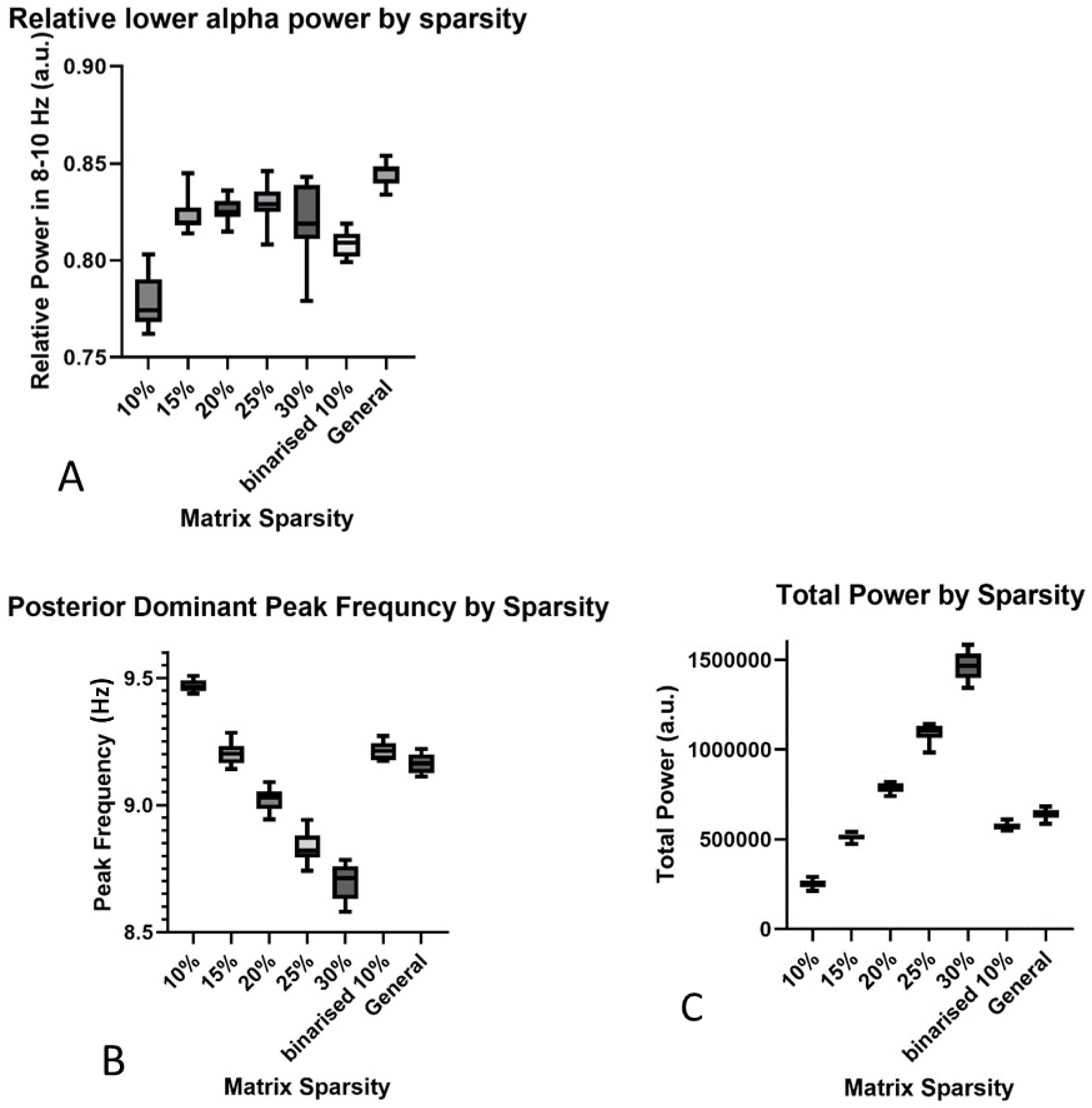
Investigating the optimal sparsity for the personalized connectivity matrices. Sparsity thresholds of 10-30 % strongest connections remaining were compared to a binarized 10% matrix and the general matrix (likewise a binarized 10% matrix). A) Relative power in the lower alpha band. B) Posterior dominant peak frequency. C) Total power. Outcome measures were averaged over the whole network and time. Due to it’s similarity in outcomes to the general model, the 15% threshold was chosen.

